# Deconstructing olfactory epithelium developmental pathways in esthesioneuroblastoma

**DOI:** 10.1101/2022.10.18.512713

**Authors:** John B. Finlay, Ralph Abi Hachem, David Jang, Nosayaba Osazuwa-Peters, Bradley J. Goldstein

## Abstract

**Importance:** Esthesioneuroblastoma (ENB) is a rare tumor arising from the olfactory cleft region of the nasal cavity. Due to the low incidence of this tumor, as well as an absence of established reagents such as cell lines or murine models, understanding the mechanisms driving ENB pathobiology has been challenging.

**Objective:** Here, we sought to apply advances from research on the human olfactory epithelial neurogenic niche, along with new biocomputational approaches, to better understand the cellular and molecular factors in low and high grade ENB and how specific transcriptomic markers may predict prognosis.

**Design, Setting, and Participants:** This was a retrospective biocomputational analysis utilizing human ENB and normal olfactory mucosal tissue and clinical outcomes data. The setting was a large academic medical center. Participants were selected based on available datasets, and included ENB patients across all four tumor grades, and patients with normal olfactory mucosa.

**Main Outcomes and Measures:** Outcomes include deconvolution analysis, differential gene expression analysis, overall survival, and immunohistochemistry. A machine learning model was used to predict cell-type proportion signatures in ENB, and Kaplan-Meier curves with associated log-rank tests were used to estimate differences in overall survival.

**Results:** We analyzed a total of 19 ENB samples (9 low grade I/II, 10 high grade III/IV) with available bulk RNA-Sequencing and survival data, along with 10 samples from normal olfactory mucosa (3 bulk RNA-Sequencing, 7 single cell RNA-Sequencing). The bulk RNA-Sequencing deconvolution model identified a significant increase in globose basal cell (GBC) and CD8 T cell identities in high grade tumors (GBC from approximately 0% to 8%, CD8 T cell from 0.7% to 2.2%), and significant decreases in mature neuronal, Bowman’s gland, and olfactory ensheathing programs, in high grade tumors (mature neuronal from 3.7% to approximately 0%, Bowman’s gland from 18.6% to 10.5%, olfactory ensheathing from 3.4% to 1.1%). Trajectory analysis identified potential regulatory pathways in proliferative ENB cells, including PRC2. Survival analysis guided by gene expression in bulk RNA-Sequencing data identified favorable prognostic markers such as SOX9, S100B, and PLP1 expression.

**Conclusions and Relevance:** Our analyses provide a basis for additional translational research on ENB management, as well as identification of potential new prognostic markers.

## INTRODUCTION

Esthesioneuroblastoma (ENB), also referred to as olfactory neuroblastoma, is a rare neoplasm that is thought to arise from the olfactory epithelium (OE) in the superior nasal cavity. ENB, first described in 1924, accounts for approximately 2-3% of sinonasal malignancies^1,2^. Given the location of the tissue of origin, local growth of ENBs can invade superiorly into the anterior skull base and brain, laterally into the orbit, or inferiorly into the inferior nasal cavity and paranasal sinuses. ENB can also metastasize to the cervical lymph nodes and distant sites. Historically, ENBs have been graded pathologically using the Hyams system, which takes into account Ki67 proliferative index and presence of necrosis^3,4^. Stage is typically determined using either the Kadish or Dulgerov system, both of which assess the extent of local primary tumor extension to the paranasal sinuses or cranial cavity, and the presence of local metastasis^5,6^. Patients can present with epistaxis, nasal congestion, olfactory loss, visual changes, headache, and in advanced disease, weight loss.

The standard of care for ENB, which has remained relatively unchanged for decades, is surgical resection followed by radiation therapy^7,8^. Chemotherapy has often been added for advanced disease^9^. Even with aggressive surgical and medical management, the 2-year survival rate of high grade ENBs remains steady at approximately 40%, and late recurrences can occur^10^.

Because of their extremely low incidence of 0.4 in 1 million, large detailed studies of ENB have been a challenge, although case series and systematic reviews have provided insights^11,12^. No reliable mouse models or patient-derived cell lines exist, making it difficult to experimentally manipulate tumor biology. In addition, the precise mechanisms driving proliferation and differentiation in the adult OE, which actively produces new neurons throughout life from resident basal stem cells, remain an area of active research. Thus, it has been difficult to contextualize the neoplastic processes driving ENB growth with regard to the specific cell identities of the OE. Due to these limitations, the use of targeted and/or immune modulating therapies has not been established in the treatment for ENB, in contrast to other cancers.

Recent advances in human olfactory stem cell biology offer a new lens through which to study ENB^13^. The adult OE is a dynamic neurogenic niche that allows for constant production of sensory neurons, sustentacular support cells, microvillar cells, and glands throughout life^14–17^. The embryonic origin of the OE is largely derived from a cranial sensory ectodermal placode, while the underlying lamina propria, housing glands, vessels and olfactory ensheathing glia cells, arises from cranial neural crest^18^. However, mouse genetic lineage tracing studies identified evidence for intermixing of placode and neural crest, reflecting the complexities of the origins of the peripheral olfactory structures in mammals^18^. In adults, a reserve population of quiescent stem cells, termed horizontal basal cells (HBCs), lines the basal lamina of the OE, and can become activated by epithelial damage to generate proliferative globose basal cells (GBCs). GBCs are heterogenous and function as amplifying neuronal progenitors or immediate neuronal precursor (INP) cells that differentiate rapidly into immature olfactory sensory neurons (iOSNs) or microvillar cells; there is also a direct lineage relationship between GBCs and submucosal Bowman’s glands.

It has been suggested that high grade ENBs arise from proliferative basal cells, with less aggressive tumors having more neuronal characteristics^19^. However, this does not take into account the diversity of differentiation trajectories and proliferative cell states within the normal OE. The expression of glandular proteins within ENB has also been reported, suggesting that tumors may be more heterogenous than originally thought^20^. An olfactory neuronal lineage origin for ENB has been supported strongly by identification of neuronal antigens such as neurofilament in the tumors^21^, or by identification of GBC-specific transcription factors such as ASCL1^22^. More recent molecular analyses have also suggested the expression of neural crest transcripts using bulk RNA-sequencing^19^, although this approach cannot provide cell type-specific resolution. As such, the cellular and molecular mechanisms driving ENB growth remain incompletely understood.

Here, we take advantage of an integrated single cell transcriptomic atlas of the human olfactory mucosa, which we previously generated using human biopsy samples, to deconvolute bulk RNA-Seq expression patterns in 19 publicly available ENB samples and 3 control samples. Our findings indicate that GBCs are significantly enriched in higher grade tumors, as expected. Differentiation trajectory analysis in normal OE cells reveals that components of the Polycomb repressive complex 2 (PRC2), a conserved epigenetic regulatory structure, are enriched in both GBC and iOSN states. We perform here immunostaining confirming that PRC2 proteins are associated with proliferative cells in ENB, even in low grade tumors. We find that lower grade tumors are enriched for olfactory ensheathing cell and Bowman’s gland transcripts, and that expression of canonical markers for these cell identities is positively correlated with overall survival. Guided by perspectives from adult olfactory cell biology, this analysis provides novel insights into factors active in ENB tumors, suggesting important mechanisms contributing to tumor growth and outcome, as well as pathologic markers for prognosis.

## RESULTS

### Pseudo deconvolution of normal OE and ENB bulk RNA-Seq samples

Given that ENBs are rare tumors, it is not surprising that single-cell transcriptomic datasets that encompass the diversity of low to high grade tumors are not yet available. Therefore, we sought to take advantage of recent innovations in transcriptomic biocomputational analyses by deconvoluting cell type signatures in available bulk RNA-Seq ENB datasets^19^, using a robust single cell reference atlas from normal human olfactory mucosa. Traditionally, bulk RNA-Seq deconvolution involves training a model with a single cell (sc) RNA-Seq dataset from the same tissue type. The model learns patterns of aggregate gene expression across different cell identities, and then analyzes bulk RNA-Seq datasets for similar expression patterns (Fig 1). The output is a relative proportion of each cell identity (as annotated in the scRNA-Seq reference dataset) within a given bulk RNA-Seq sample. Here we refer to our analysis as pseudo deconvolution, since the scRNA-Seq training dataset is from normal olfactory mucosa cells, not ENB. This is highly useful, since the deconvolution can identify patterns of gene expression within ENBs that map directly to specific cell states within the tissue from which tumors arise. As a control, we deconvoluted three bulk RNA-Seq samples obtained from normal human OE.

**Figure 1:**
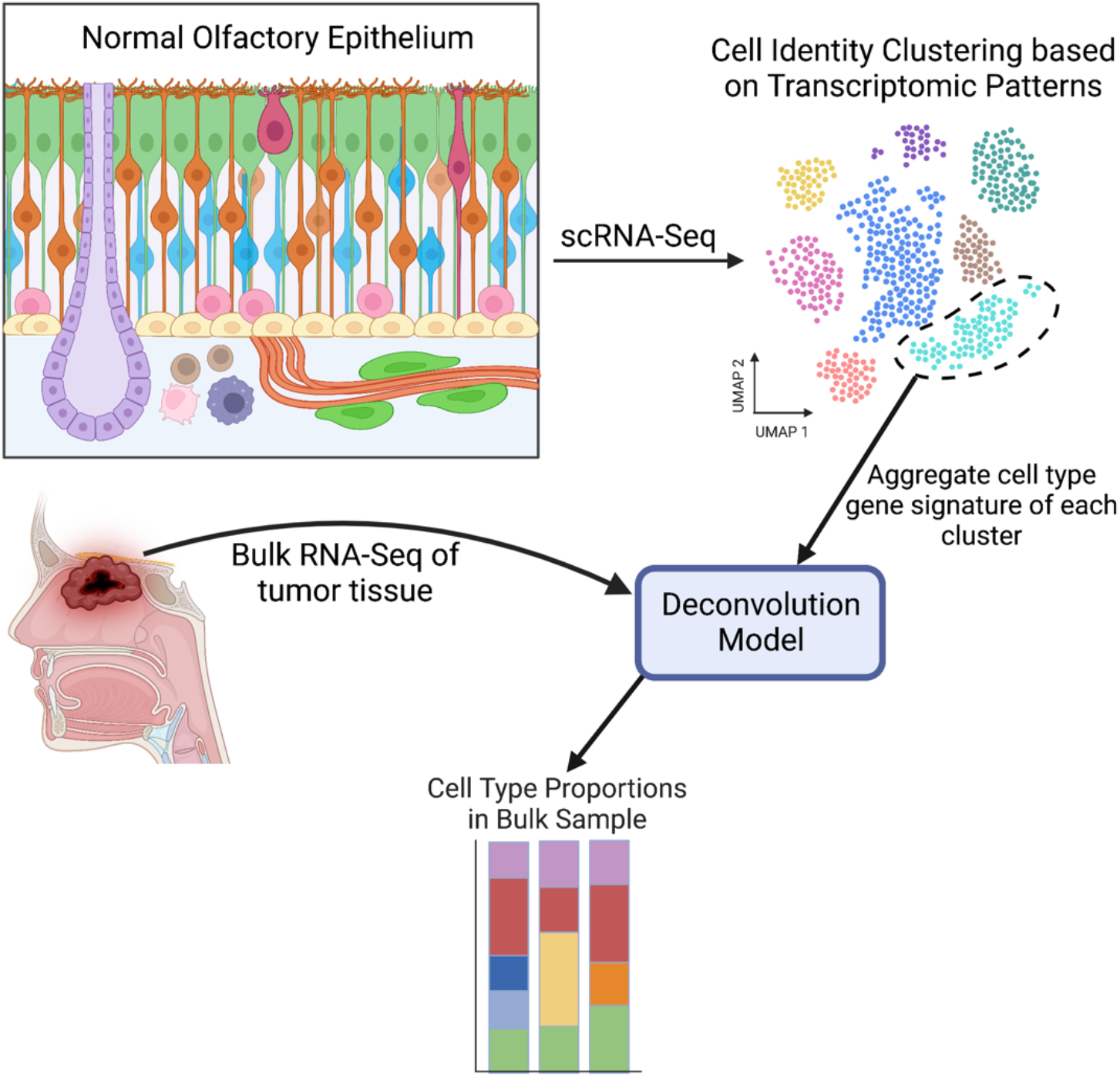
Schematic of the deconvolution workflow. Diagram of normal olfactory epithelium in the top left colored by cell identity (HBC: yellow; GBC: pink; immature OSN: blue; mature OSN: orange; Sustentacular: green; Bowman’s Gland: purple; Olfactory ensheating cell:lime green in lamina propria; immune cells: round cells in lamina propria). The schematic depicts the two main inputs into the deconvolution model: a reference single-cell RNA-Seq dataset along with bulk RNA-Seq data. The output is a stacked bar graph with estimated proportions of cell identity per bulk RNA-Seq sample.

Across the three normal OE samples, we observed similar patterns of cell type abundance, with approximately 10% neural lineage cells, 10% immune cells, 20% Bowman’s glands, and a significant proportion mapping to respiratory cell types, which is expected given that the OE is generally interspersed with patches of respiratory epithelium in humans (Fig. 2A). When we explored deconvolution of low and high grade ENB samples, we identified a clear enrichment of OE cell types in ENB (Fig. 2A). Specifically, there is a significantly greater proportion of GBCs in high grade ENB compared to normal OE, and significantly fewer mature OSNs, as compared to low grade tumors (Fig. 2B). Note, GBCs generally represent approximately 0.25% of cells captured in our normal adult olfactory mucosa biopsies, so they artificially appear absent from deconvolution of bulk seq samples given the contribution of their unique transcripts to overall reads. Sustentacular and microvillar cells, which can both arise from GBCs, were either decreased (sustentacular) or remained the same (microvillar) in ENBs, when compared to normal OE. Interestingly, Bowman’s glands, which were found to be relatively abundant (20% proportion) in normal olfactory mucosa biopsies, trended down with higher tumor grade, with significantly fewer Bowman’s gland gene signatures in high grade ENBs (Fig. 2B). Similarly, compared to low grade tumors, high grade ENBs had significantly fewer olfactory ensheathing cells, a neural crest-derived glial cell similar to non-myelinating Schwann cells. We did not observe a significant difference in proportions of proliferative respiratory suprabasal cells, confirming that the model uses more than a simple panel of proliferation genes to map tumor cell identity. Finally, there have been conflicting data on the existence of a positive correlation between tumor grade and degree of T cell infiltration in ENB^19,23,24^. With our model, we find that higher grade ENBs do have a higher proportion of CD8 T cells.

**Figure 2:**
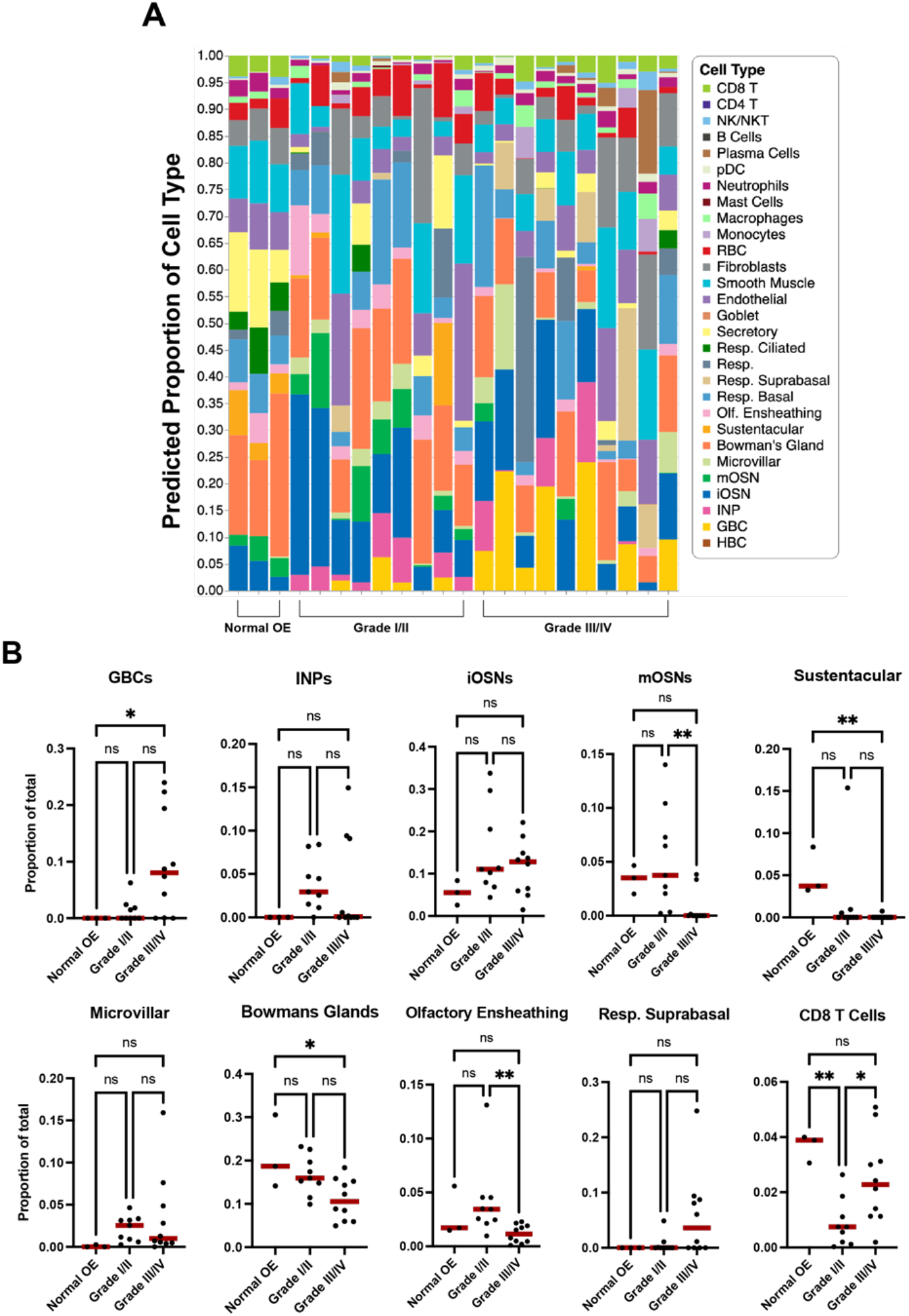
Pseudo-deconvolution of ENB bulk RNA-Seq defines proportions of cell type-specific transcriptomics across low and high grade tumors. **A**) Stacked bar chart depicts predicted proportions of cell types in bulk tumors when mapped to control adult human olfactory mucosa using the deconvolution model. **B)** Statistical differences in selected ENB cellular composition, based on one-way ANOVA followed by post-hoc Tukey’s test (* p<0.05, ** p<0.01).

These data suggest that while lower grade ENBs do contain more differentiated neural components, they may also have significant proportions of other differentiated cell-type lineages including Bowman’s glands and olfactory ensheathing cells. It is also interesting to note that higher grade tumors do not map entirely to rapidly proliferating basal cells, and that they still contain significant proportions of neuronal cell type signatures. While the available bulk RNA-seq data sets are felt to contain predominantly tumor cells, it is impossible to exclude some contribution of normal overlying surface epithelium, either olfactory or respiratory, which could contribute to overall transcripts.

### Trajectory analysis in normal OE samples reveals potential regulators of tumorigenesis

Given the olfactory cell lineages identified in both low and high grade ENBs, we hypothesized that regulatory mechanisms controlling normal olfactory proliferation and/or differentiation are likely to play a role in ENB tumorigenesis. One likely candidate is the polycomb family. PRC2 is an epigenetic modifier that regulates transcription in developing and adult stem cell niches via modulation of chromatin structure. We have previously identified expression of core PRC2 proteins, including EZH2, EED, and SUZ12, in OE sensory lineages, and EZH2 histone methyltransferase activity appears to regulate murine GBC proliferation^25,26^. Therefore, we utilized trajectory analysis on our single cell dataset containing 7 normal human olfactory mucosal biopsies to visualize where PRC2 gene expression is up-regulated along human OE cell differentiation trajectories (Fig. 3A). We identified cellular states across three different lineage trajectories and used previously identified cell type-specific markers to confirm cell identity^13^ (Fig. 3B). Specifically, we identified a cluster of HBCs (TP63^+^, KRT5^+^) as the starting point of the differentiation trajectory. From here, trajectories could either progress along pseudo-time toward a fully mature neuron (mOSN, RTP1^+^, GNG13^+^), a microvillar cell (ASCL3^+^), or a sustentacular cell cluster (SOX2^+^, ERMN^+^). Intervening GBC and immediate neuronal precursor (INP) stages were also identified.

**Figure 3:**
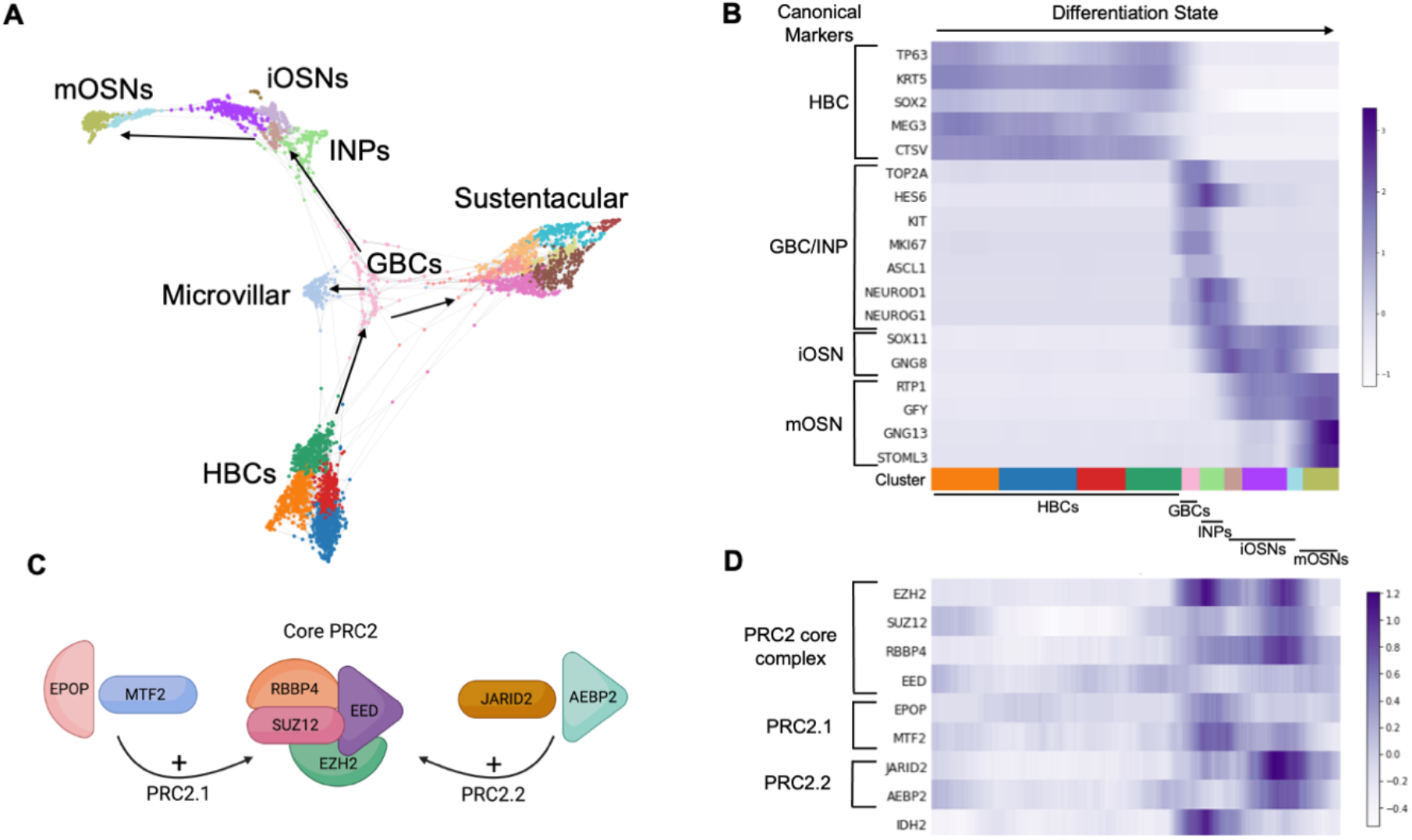
Patterns of gene expression in the normal differentiating olfactory epithelium. **A)** Trajectory UMAP depicting differentiation states across pseudo-time, in 7 normal olfactory mucosa biopsies scRNA-seq from Durante et al., 2020 and Oliva et al., 2021^13,38^.**B)** Heatmap showing canonical cell type-specific markers for selected OE cell stages. **C)** Schematic showing that the core PRC2 protein complex can combine with either EPOP+MTF2 to form PRC2.1 or with JARID2+AEBP2 to form PRC2.2. **D)** Heatmap showing PRC2 gene expression (and IDH2). (HBC: horizontal basal cell; GBC: globose basal cell; INP: immediate neuronal precursor; iOSN: immature olfactory sensory neuron; mOSN: mature olfactory sensory neuron; PRC2: Polycomb repressive complex 2).

Having established an *in silico* model of human OE differentiation, we next focused on transcripts for core PRC2 components (EZH2, SUZ12, EED, RBBP4) or subunits specific to PRC2.1 (MTF/PCLs, EPOP) or PRC2.2 (JARID2, AEBP2) complexes (Fig. 3C)^27^. Distinct PRC2 complexes with unique subunit composition confer different transcriptional regulation to certain cell phenotypes in other self-renewing systems, such as the skin or hematopoietic system^28^; however, these potential roles in the adult OE remain to be defined. As a cell differentiates from an HBC into a mature OSN, we observed a pattern of significantly increased expression in PRC2 transcripts in GBC cell identities (Fig. 3D). Of note, this first peak is associated with a single peak in EPOP and MTF2, two constituents of PRC2.1. As cells continue on to INPs, there is a minor decrease in many of these genes, followed by a second peak in expression prior to an iOSN maturing into an mOSN; this second peak, in contrast to the first, corresponds with a single peak in JARID2 and AEBP2, constituents of PRC2.2. Once a cell is a fully mature neuron, there is consistently low expression of PRC2 transcripts. Unlike in the neuronal trajectory, we did *not* detect differences in expression of Polycomb subunits along the sustentacular differentiation trajectory, consistent with a neuron lineage-specific function for PRC2.

We also queried expression of IDH2 along these trajectories, given that mutations in IDH2 have been one of the only recurrent mutations reported in high grade ENB^19^. We find that like PRC2 genes, IDH2 expression peaks at the GBC to INP cell identity (Fig. 3D). This finding serves as further confirmation that higher grade ENBs are likely expressing cellular programs that most closely resemble those of GBCs. Furthermore, our trajectory analysis reveals that PRC2 genes are selectively expressed in the GBC/iOSN cellular states in human OE.

### PRC2 is associated with proliferation in ENB

Because PRC2 is active in proliferative neural progenitors in the mouse^25,26^ and normal human OE, we asked if there was any link between PRC2 expression and proliferation in ENB. Using the previously published datasets of bulk RNA-Seq from ENBs and normal olfactory mucosa, we divided samples into normal OE, Hyams Grade I/II ENB, and Hyams Grade III/IV ENB, as in Fig. 1C. Higher grade tumors have a greater percentage of Ki67^+^ cells, reflective of increased proliferative activity. We compared expression of PRC2 transcripts (EZH2, EED, SUZ12, RBBP4) across these three groups and found a significant increase in expression of EZH2, EED, and SUZ12 in the higher grade tumors compared to lower grade (Fig. 4A). Because we observed that accessory proteins associated with PRC2.1 (EPOP, MTF2) peaked at the GBC/INP state and those associated with PRC2.2 (JARID2, AEBP2) peaked at the iOSN state, we asked if there were detectable differences across ENB grade. Interestingly, there is a significant increase in EPOP expression in high grade compared to low grade tumors, with no differences between tumor grade in PRC2.2 protein expression (Fig. 4B). This suggests that PRC2.1 specifically may play a role in ENB malignancy.

**Figure 4:**
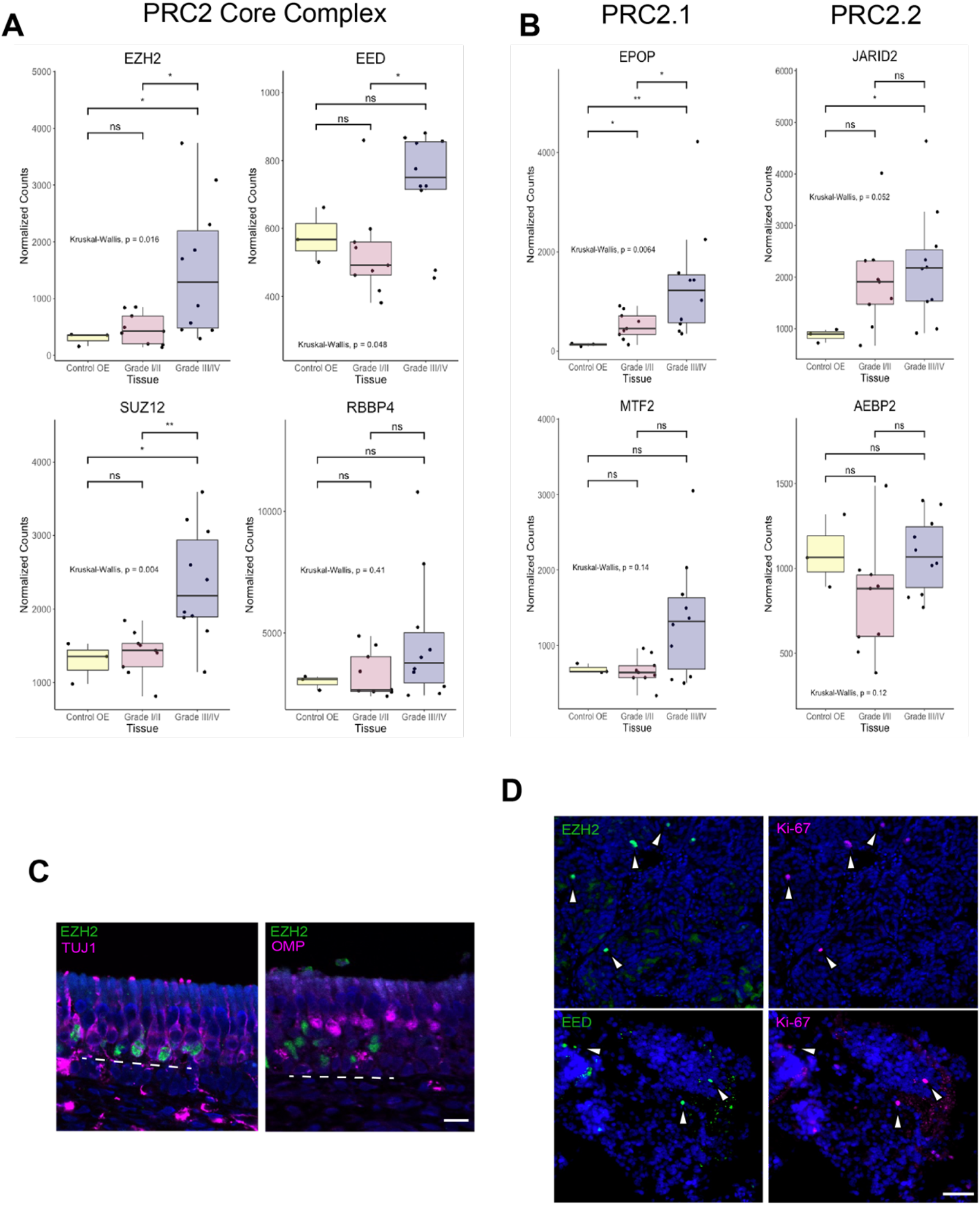
PRC2 complex expression in ENB. **A)** PRC2 transcripts in the core complex are mostly up-regulated in high grade ENB. **B)** EPOP expression is significantly increased in high grade ENB compared to low grade, with no differences between low and high grade in proteins that form the PRC2.2 complex. For A and B, statistical significance assessed by Kruskal-Wallis test with post-hoc Dunn test for multiple comparisons (* p<0.05, **p<0.01). **C)** EZH2 is present in basal cells and immature neurons (TUJ1^+^) but not in mature neurons (OMP^+^) in the normal human OE. **D)** Immunohistochemically, EZH2 and EED co-localize with Ki67 in a grade I ENB. Scale bars in C and D are 50μm.

Next, we used immunohistochemistry (IHC) to stain for PRC2 proteins. We first confirmed presence of EZH2 protein in the normal human OE, and validated that its expression is limited to basal cells and immature neurons (TUJ1^+^) (Fig. 4C). We then examined PRC2 protein expression in an ENB sample and co-stained with anti-Ki67 to assess proliferation. Although our staining was performed on a Grade I tumor, Ki67^+^ cells were identified throughout the sample (Fig. 4D). Staining revealed that nearly all Ki67^+^ cells were also EZH2^+^. Similarly, many Ki67^+^ cells stained positive for EED. Despite expression of PRC2 in proliferative GBCs and in immature neurons in normal OE, in the tumor sections we only identified PRC2 protein expression in proliferative cells, marked by Ki67. Together these results suggest that expression of core PRC2 proteins is associated with proliferation in ENB both at the RNA and protein level.

### ENB expression of canonical Bowman’s gland or olfactory ensheathing glia transcripts correlates with favorable prognosis

With the finding that higher grade tumors have significantly decreased expression of Bowman’s gland and olfactory ensheathing cell transcriptional identities, we asked whether expression of key marker genes of these cell types was associated with favorable prognosis. Differential expression analysis in our single-cell atlas of 7 normal human olfactory mucosa samples identifies PLP1 and S100B as genes unique to olfactory ensheathing cells, while high levels of SOX9 is specific to Bowman’s glands. SOX9 expression is, however, also present in the subset of Trpm5^+^ microvillar cells (Fig. 5A). To assess impact on prognosis, we queried the ENB bulk RNA-Seq dataset and divided tumors into low or high expression groups based on whether expression of a given gene fell below or above the median expression of the gene across all tumors. Then, we generated Kaplan-Meier plots and ran log-rank tests to determine statistical significance. Of particular interest, we found that tumors with higher expression of PLP1, S100B, and SOX9 all confer a statistically significant increase in survival (Fig. 5B). Because TRPM5^+^ microvillar cells also express SOX9, we ran a similar analysis with TRPM5 expression but did not observe any differences in survival, suggesting that the finding are related to SOX9^+^ Bowman’s gland cells, and not microvillar cells (Fig. 5B).

**Figure 5:**
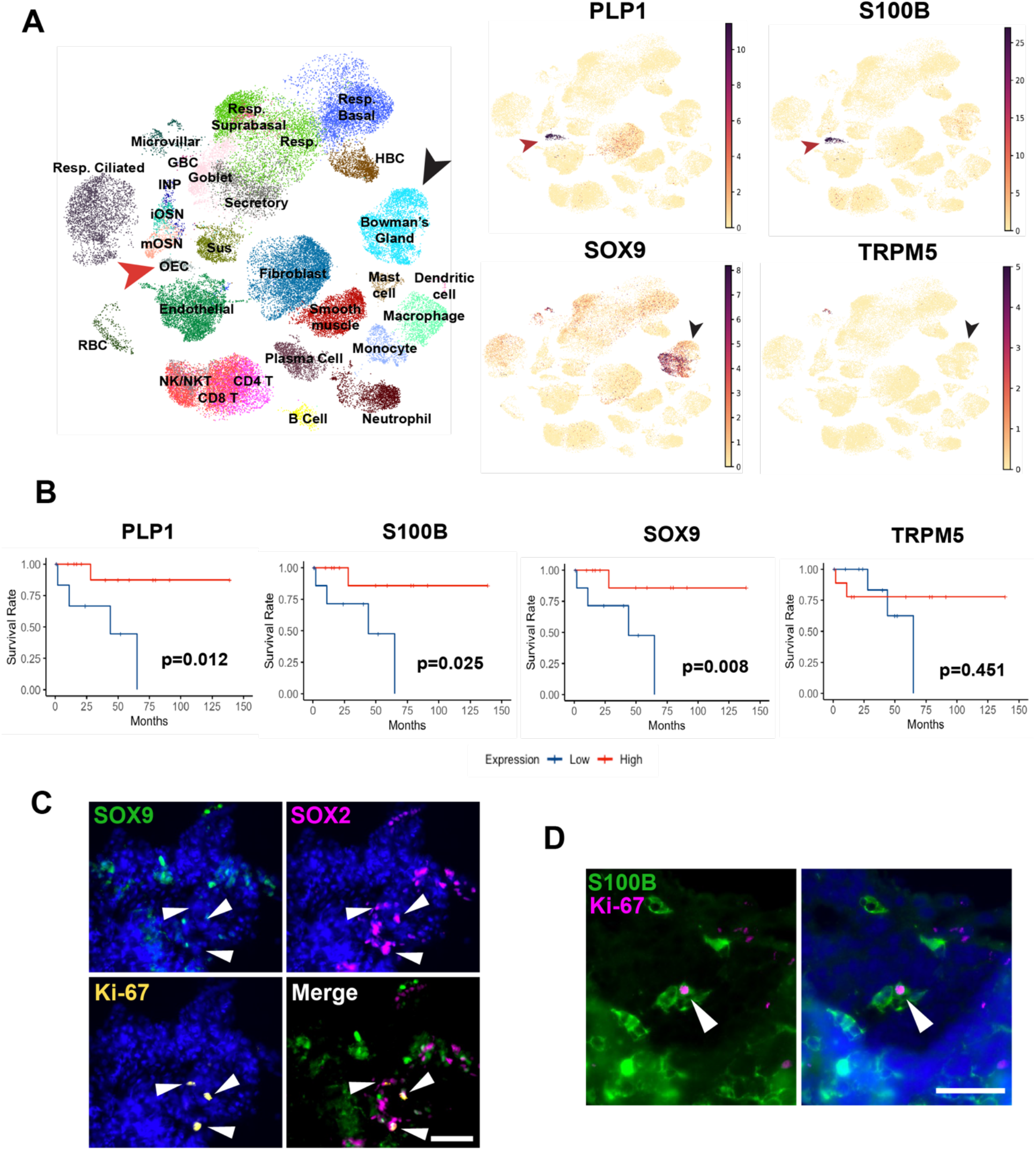
Glandular and olfactory ensheathing cell markers are present in low grade ENB and are associated with increased survival. **A)** UMAP visualization of non-tumor olfactory mucosa scRNA-seq data depicts the cell typespecific expression of selected transcripts. PLP1 and S100B are expressed in olfactory ensheathing cells (OEC, red arrow); SOX9 is highly expressed in Bowman’s glands (BG, black arrow) and the TRPM5^+^ subset of microvillar cells. Scale bar shows normalized counts. **B)** Kaplan-Meier survival curves showing significant survival advantage in ENB tumors with higher expression of BG and OEC markers. Statistical significance determined by log-rank test. **C)** SOX9^+^ cells are rarely Ki-67^+^ and do not co-localize with SOX2. In contrast, some SOX2^+^ cells show evidence of proliferation based on Ki-67 staining (arrowheads). **D)** The presence of S100B^+^ cells in a grade I ENB. Ki-67 co-staining confirms the presence of the occasional proliferative S100B^+^ cell (arrowhead). In A, resp = respiratory; RBC = red blood cell; sus = sustentacular; in C and D, scale bar = 50 μm.

As controls, we assessed the effects of Hyams grade, Dulgeurov T stage, and MKI67 expression (gene for Ki67 protein) on survival in these data sets. Hyams Grade and Dulgeurov T stage were both approaching but did not achieve statistical significance, while higher MKI67 expression was associated with significantly worse survival (Fig. S1). Our analysis suggests that, in this ENB dataset, PLP1, S100B, and SOX9 expression levels are all better indicators of survival than the standard grading and staging systems.

For further validation of these prognostic markers, we analyzed S100B and SOX9 protein in an ENB tumor by immunohistochemistry. Our staining confirms the expression of nuclear SOX9 or SOX2 protein in ENB, and these transcription factors are mutually exclusive (Fig. 5C). Also, we did not identify co-expression of Ki67 in SOX9^+^ cells. We also confirmed the expression of S100B protein in ENB and do identify occasional co-expression of Ki67 in S100B^+^ cells (Fig. 5D). Of interest, in normal adult human olfactory mucosa, S100B expression is highly specific to olfactory ensheathing glia, as it is in rodent^29,30^. Although prior reports have referred to S100B in ENB as a marker for sustentacular cells^31^, we are not aware of any evidence in mouse or human that this protein is present in sustentacular cells (Fig. 5A). We conclude that S100B expression in ENB is likely reflective of the presence of ensheathing glia, or possibly other neural crest-derived elements related embryonically to the surrounding mesenchyme.

## DISCUSSION

The findings presented here have several implications regarding mechanisms of ENB tumor growth and prognosis. By leveraging new insights from the adult human olfactory epithelial stem cell niche and new biocomputational techniques, we sought to better understand the biology of a rare and clinically challenging olfactory cleft neoplasm.

Several challenges exist when considering ENB treatment. At present, no accepted or standard targeted / biologic therapy for ENB has been identified. Therefore, higher grade tumors have been managed with conventional regimens often using cisplatin and etoposide, or cyclophosphamide/vincristine, along with surgery and radiation^2,32^. Significant potential morbidity must be considered with achieving widely negative margins of surgical resection, due to anatomy of the olfactory cleft and adjacent structures. Morbidity can involve need for resection of orbit or intracranial contents. Furthermore, bilateral anterior skull base resection and removal of olfactory bulbs renders patients permanently and irreversibly anosmic. Thus, the identification of targets for biologic or novel therapies, with potential to be effective in advanced ENB, is needed.

The findings presented here provide a rationale for potential treatment strategies. Leveraging advances in our understanding of the mechanisms regulating olfactory neurogenesis and, specifically, GBC proliferation and differentiation, and mapping ENB transcriptional profiles to normal adult human olfactory data sets at single cell resolution, we identify new insights regarding ENB. A key finding involves a likely role for PRC2 in proliferative ENB cells. It is intriguing to consider that the enzymatically active PRC2 subunit, EZH1/2, is a druggable target^33^. Indeed, PRC2 is a treatment focus for several malignancies, including hematologic and solid tumors^34,35^. In considering drug development, there is also a focus on potential PRC2-independent roles for EZH2 in cancer^36^.

In addition to potential utility in cases of ENB metastasis or recurrence, a targeted ENB therapy may be useful as a neoadjuvant drug for bulky ENB tumors, to permit removal via a hemi-skull base resection that could spare the contralateral olfactory structures and thus preserve olfaction. Our results provide a basis for further research regarding targeted therapy for ENB.

Another novel finding from our analysis is the potential utility of specific prognostic markers in ENB. Specifically, we found here that expression of the Bowman’s gland transcription factor SOX9, and also the olfactory ensheathing glia markers S100B or PLP1, all correlate significantly with improved prognosis in this data set including 19 ENB subjects. Considering the low incidence of ENB, our approach mapping existing transcriptional data sets to robust normal human olfactory single cell data provided a means to identify cell type-specific transcripts of interest for survival data query.

Together, our findings suggest important mechanisms contributing to ENB tumor growth, as well as pathologic markers for prognosis. We anticipate that these insights will be useful in guiding further translational studies. More broadly, these mechanistic insights will be helpful in considering development of future personalized approaches to the management of this rare and challenging tumor.

## MATERIALS AND METHODS

### Bioinformatics

#### Pre-processing of bulk RNA-Seq samples

Bulk RNA-Seq datasets were queried from the NCBI Gene Expression Omnibus from GSE118995 (19 ENB samples)^19^, and GSE80249 (3 normal OE samples)^37^. FASTQ files for each patient sample were downloaded and imported into Galaxy (v22.05.1) via SRA for processing. Low quality sequencing reads were trimmed from paired-end data using Trimmomatic (v0.38.0) with default settings and no Illumina barcode trimming. Trimmed datasets were then aligned to hg19 using HISAT2 (v2.2.1) with default settings. Counts per gene were calculated with featureCounts (v2.0.1) with “Count fragments instead of reads” enabled under “options for paired-end reads,” and all other default settings. Finally, annotateMyIDs (v3.14.0) was used to link Entrez IDs to Gene names.

#### Bulk RNA-Seq deconvolution

Deconvolution of bulk RNA-Seq datasets with an integrated single cell RNA-Seq reference dataset was performed in Python (v3.9.0) using Scanpy (v1.9.1). Single cell RNA-Seq datasets used as a reference in the deconvolution model were imported from GSE139522^13^ and GSE184117^38^. Specifically, we used all 4 patients from Durante et al., 2020 and the 3 normosmic patients from Oliva et al., 2021. Datasets were processed, integrated using scvi-tools (v0.17.4), and annotated by cell type cluster, as described previously^39^.

For deconvolution, we utilized RNA-Sieve (v0.1.4)^40^. Once an integrated scRNA-Seq dataset was properly annotated by cell type, we subset this to include only the 4 olfactory mucosal biopsies from Durante et al., 2020 in order to streamline computational workflow. A concatenated counts matrix with counts from all 19 ENB and 3 normal OE bulk RNA-Seq dataset was imported. We eliminated genes only present in either the scRNA-Seq dataset or the bulk RNA-Seq datasets using index.intersection(). Raw counts for scRNA-Seq and bulk RNA-Seq datasets were prepped for running the deconvolution model with default settings, as described^40^. The deconvolution model was trained by running model_from_raw_counts() on the processed scRNA-Seq reference dataset. Then the model was run with model.predict(), using default settings. Output graphs were produced using Altair (v4.2.0). Statistical comparisons between predicted cell-type proportions were run in Graphpad Prism 9.

#### Trajectory Analysis and UMAPs

Trajectory analysis was performed using Scanpy (v1.9.1) and the integrated scRNA-Seq dataset with 7 normal human olfactory mucosa biopsies, described above. First, we subset out just olfactory epithelial clusters (HBCs, GBCs, INPs, iOSN, mOSN, Microvillar, and Sus), excluding Bowman’s glands. Data were normalized with sc.pp.normalize() with a target_sum of 1e4. Next, sc.tl.pca() was run, followed by sc.pp.neighbors() with 5 neighbors and 30 principal components. A plot was generated with sc.tl.draw_graph() followed by sc.pl.draw_graph(), with default settings. We performed de-noising by running sc.tl.diffmap() followed by sc.pp.neighbors() with 5 neighbors. Data were re-clustered with sc.tl.leiden with a resolution of 2.0. Then partition-based graph abstraction was run with sc.tl.paga() followed by sc.pl.paga(), then sc.tol.draw_graph(), and finally sc.pl.draw_graph(), all with default settings. Cell identities were analyzed with feature plots. Small clusters of cells that did not align with any of the known cell markers were filtered out. The process was re-run iteratively beginning with sc.tl.pca() for a total of three times to ensure purity of cell identities within the trajectory plot. To produce the trajectory marker heatmap, data were logarithmized and then scaled with sc.pp.log1p() and sc.pp.scale() in order to allow for visualization of variably expressed genes on one plot. Data were plotted with sc.pl.paga_path(), using default settings. All UMAPs and featureplots were produced with normalized count matrices and plotted using sc.pl.umap().

#### Down-stream analysis of RNA-Seq samples

Concatenated bulk RNA-Seq count matrices (19 ENB specimens, 3 normal OE), were loaded into R (v4.1.1) for further differential expression analysis. To compare gene expression between tumor grades, DESeq2 (v1.32.0) was run, using default settings. Counts were normalized with counts(dds, normalized=T). Box plots of counts were produced with plotCounts(), and stat_compare_means() was run to perform Kruskal-Wallis testing among all three groups, followed by post-hoc Dunn test for multiple comparisons.

Survival (v3.3-1) and survminer (v0.4.9) were used to perform survival curve analysis based on gene expression. We first created a matrix with survival data from each of the 19 ENB patients using available data from Classe et al., 2018^19^. A DESeq2 object using only the 19 ENB bulk RNA-Seq count matrices was produced as described above, with normalized counts. For a given gene, we then calculated the median expression among all 19 tumors. We divided the dataset into a “low” and “high” expression group using approximately the median value as a cutoff. Then, a survival model integrating the combined count matrix with the survival outcomes matrix was created using survfit(), with overall survival as the outcome measure. A log-rank test p-value was calculated using surv_pvalue() and the Kaplan-Meier graph was plotted with ggsurvplot().

### Immunohistochemistry

Normal olfactory mucosal and ENB biopsies were collected in the operating room on ice in Hanks’ Balanced Salt Solution (HBSS, Gibco) +10% FBS. Surgical specimens were washed twice in phosphate buffered saline (PBS) and fixed for approximately 3 hours in 4% paraformaldehyde (Sigma, St. Louis). Tissue was washed two times in PBS and incubated at 4°C for 3-5 days in 30% sucrose, 250mM EDTA, and PBS. Specimens were flash frozen with liquid nitrogen in OCT (VWR, Radnor, PA), cut into 10μm thick sections using a CryoStar NX50 cryostat (Thermofisher), and collected on Superfrost plus glass slides (Thermofisher).

Tissue sections on slides were incubated in PBS for 5 minutes, followed by dehydration-rehydration in ethanol (1 min 70% ethanol, 1 min 95% ethanol, 1 min 100% ethanol, 1 min 95% ethanol, 1 min 70% ethanol, 5 min PBS). Antigen retrieval was performed using citrate-based unmasking solution (Vector Laboratories, Newark, CA) by steaming for 45 minutes. Tissue was washed once in PBS for 5 minutes. Sections were blocked with blocking buffer (5% normal goat serum, 0.1% TritonX-100 in PBS) for 1 hour in a moist chamber at room temperature. Tissue was incubated in primary antibody overnight at 4°C in a moist chamber diluted in blocking buffer with the following dilutions: Anti-Tubulin β3 (BioLegend, clone TUJ1, 1:500); Anti-OMP (Santa Cruz, sc-365818, 1:500); Anti-EZH2 (Abcam, clone 11/EZH2, 1:500); Anti-EED (Cell Signaling Technologies, clone E4L6E, 1:50); Anti-SOX2 (eBioscience, 14-9811, 1:50); Anti-SOX9 (Cell Signaling Technologies, clone D8G8H, 1:50); S100B (Abcam, clone 4C4.9, 1:100); Anti-Ki-67 mouse (DAKO, clone MIB-1, 1:50); Anti-Ki-67 rabbit (Abcam, ab15580, 1:250). Sections were washed 3 times for 5 minutes each in PBS+0.3% TritonX-100 and incubated with appropriate fluorescent conjugated secondary antibody (Jackson ImmunoResearch, West Grove, PA) for 45 minutes at room temperature. For EZH2 and SOX9 antibodies, in place of fluorescent conjugated secondary antibody incubation, a TSA fluorescein system (Akoya Biosciences, Marlborough, MA) was used to amplify signal. Tissue was washed 3 times for 5 minutes each in PBS+0.3% TritonX-100, incubated for 3 minutes in Hoechst nuclear stain (1:1000 in H2O, Thermofisher), and mounted with Vectashield (VectorLaboratories, Burlingame, CA). Images were acquired using a Leica DMi8 microscope system, and Leica Application Suite X software (v3.7.5). Images were processed in ImageJ (v2.3.0) and scale bars were drawn using metadata in the .lif image files.

### Statistics

All statistical tests were performed in R (v4.1.1) or Graphpad Prism 9. Datasets were assessed for normal distribution, and the appropriate follow-up test (ANOVA vs. non-parametric) was performed.

## Data Availability

All datasets used in this manuscript are publicly available on GEO, with accession numbers listed above. Code used in analysis and to produce figures will be deposited on Github upon final acceptance of the manuscript.

## Study Approval

Research was performed under approved protocol from Institutional Review Board at Duke University, #Pro00088414.

## Acknowledgements

We thank all of the patients who played a role in making this study possible. Graphical schematics in this manuscript were created with BioRender.

## Author Contributions

Conceptualization: J.B.F., B.J.G.; Methodology: J.B.F., B.J.G.; Data analysis: J.B.F., B.J.G., N.O-P.; Writing or manuscript review: J.B.F., R.A-H., D.J., N.O-P., B.J.G.; Funding acquisition: B.J.G.

---

**Figure S1:**
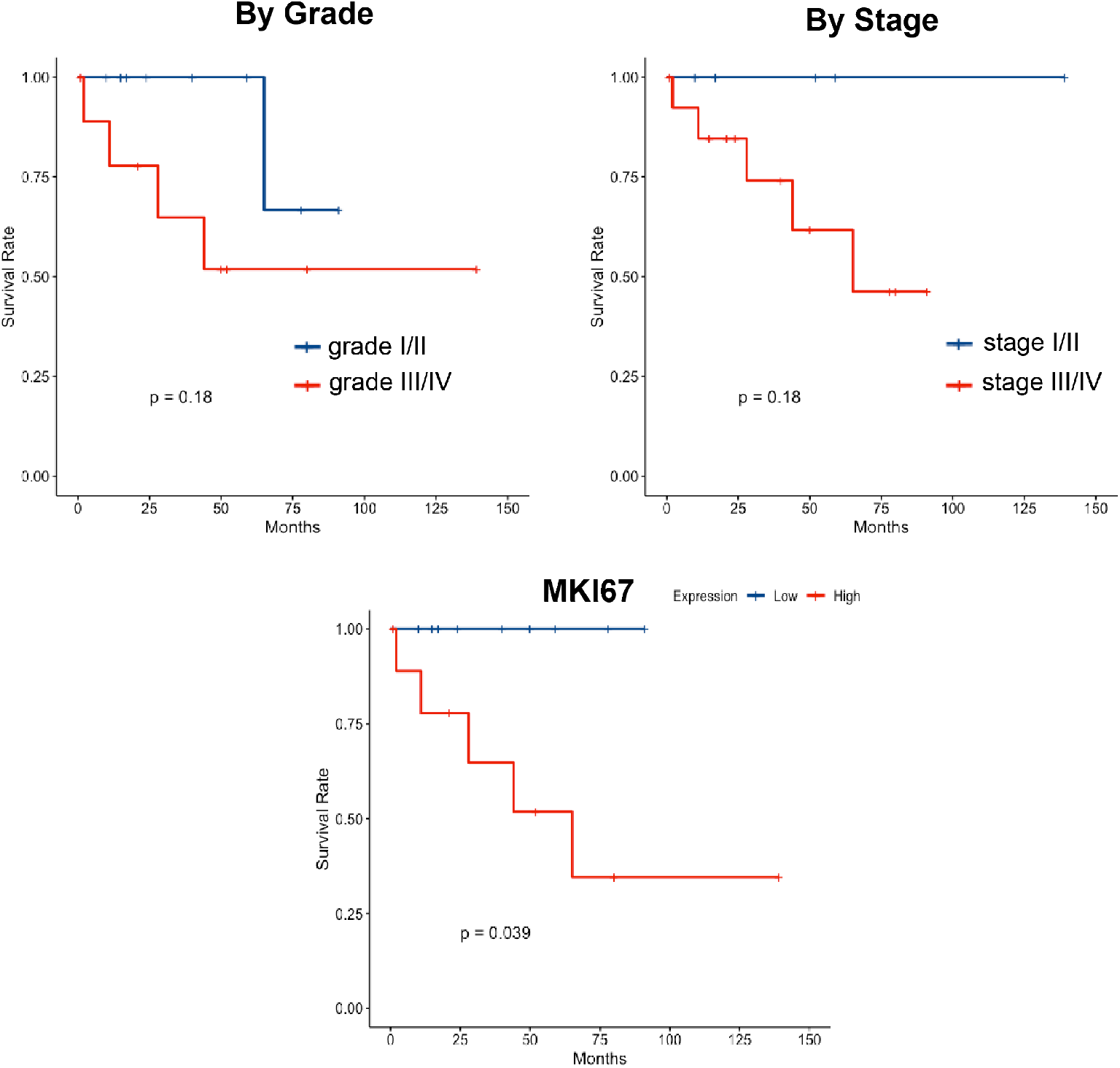
Additional Kaplan-Meier survival curves based on Hyams Grade, Dulguerov T-stage, and MKI67 gene expression. Log-rank test used to determine statistical significance.

